# The value of volunteer surveillance for the early detection of biological invaders

**DOI:** 10.1101/2022.06.24.497568

**Authors:** Frank van den Bosch, Neil McRoberts, Yoann Bourhish, Stephen Parnell, Kirsty L. Hassall

## Abstract

Early detection of invaders requires finding small numbers of individuals across large landscapes. It has been argued that the only feasible way to achieve the sampling effort needed for early detection of an invader is to involve volunteer groups (citizen scientists, passive surveyors, etc.). A key concern is that volunteers may have a considerable false-positive and false-negative rate. The question then becomes whether verification of a report from a volunteer is worth the effort. This question is the topic of this paper.

We show that the maximum plausible incidence when the expert samples on its own, 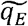, and the maximum plausible incidence when the expert only verifies cases reported by the volunteer surveyor to be infected, 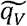, are related as

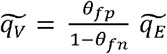

Where θ_fp_ and θ_fn_ are the false positive and false negative rate of the volunteer surveyor, respectively. We also show that the optimal monitoring programme consists of verifying only the cases reported by the volunteer surveyor if

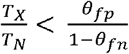

Where *T*_N_ is the cost of a sample taken by the expert and *T*_X_ is the cost of an expert verifying a case reported by a volunteer surveyor.

## INTRODUCTION

Early detection is a key requirement for successful eradication or containment of exotic invasive species [1]. Early detection requires finding small numbers of individuals across large and complex landscapes. The sampling effort and budget needed to achieve this are often well beyond the capacity of regulatory surveys. This is exemplified by first reports of invaders being made by non-specialists not involved in official regulatory surveys. For example, shot blight caused by *Siroccus tsugae* infecting Atlantic cedar (*Cedrus atlantica*) and the oriental chestnut gall wasp (*Dryocosmus kuriphilus*) were both reported first by non-specialists in the UK [2, 3]. It has been argued that the only feasible way of achieving the sampling effort needed to meet the biosecurity objective of early detection of an invader is to involve volunteer groups in data collection [4]. In response, there has been a significant increase in the number of citizen science programmes for invasive species in recent years [5].

A key concern with sightings of exotic invaders reported by volunteers is the quality of the data. It is to be expected that sightings reported by non-specialists have a considerable false-positive and false-negative rate. A study on the ability of citizen scientists to identify bumblebee species, for example, showed that, depending on the observer and bumblebee species, as few as 20-80% of the bumblebees were named correctly [6]. For a range of amphibians it was found that the false-positive rate ranged from 0.01 to 0.09 [7], considerably better than the bumblebee recognition. These results imply that a reported first sighting of an invader by a volunteer cannot be taken as conclusive proof that the invader has entered the area of interest. Verification of the sighting by an expert is essential. Conversely, the absence of detection by volunteers is not a proof of absence [8].

The question then becomes whether verification of a report from a volunteer is worth the effort or if it is more cost effective when the expert goes directly into the field themselves to sample. What is the value of volunteer reporting for the early detection of an exotic invader if the volunteer is error-prone? That question will be the topic of this paper. We will restrict our attention to pests and diseases of plants. This choice is motivated by the fact that many pests and diseases leave signs or symptoms on plants and plants do not move, enabling the expert verifying the report from the volunteer without having to deal with the additional problem of the observational unit (the plant) having moved and becoming untraceable. In the paper we will use the terminology of an infectious plant disease, but the results hold for plant pests as well.

Surveillance for early detection of invading plant pathogens often proceeds in two stages:

1. <u>Disease freedom</u>. Surveillance is started when the pathogen is believed not yet to be present. This implies that for one or more surveillance rounds no detections are made. However, since sampling is a stochastic process it might be that the pathogen is present but missed by chance. The important question is thus, what the incidence (fraction of plants infected) could be, although still missed by chance, when no detections are made.
2. <u>First detection</u>. At some point in the sequence of surveillance rounds an infected plant will be found for the first time. This establishes that the invader has arrived. The question then is whether the surveyor found the very first case or that a considerable fraction of the plants is already infected.

Eradication and containment programs are very expensive and their total cost depends on the disease incidence at the start of the management program. If initially too little resource is allocated to the eradication/containment programme the disease will escape control and the costs to get the outbreak eventually under control increase sharply. Therefore, the main interest is to know the extreme cases one may face upon first detection so that sufficient resources can be allocated. More precisely, we are interested in the upper limit of the *Z*% confidence interval of the incidence. Figure 1a illustrates this where the probability *P*, of incidence *q*, is plotted. Throughout the paper we will calculate such upper limits, 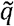, of the incidence to be expected. Throughout the paper, we refer to this as the maximum plausible incidence, with the concept of plausibility resting on our assumptions about the robustness of statistical sampling theory. This upper limit is, as shown in figure 1a, calculated from

**Figure 1:**
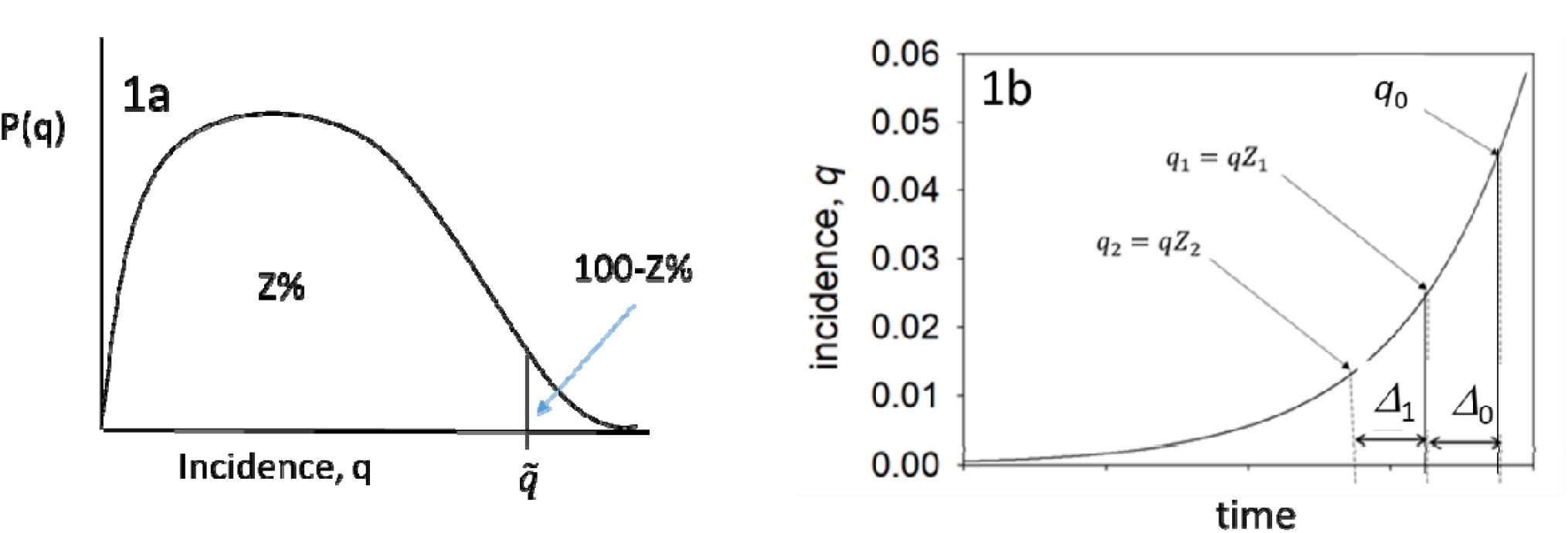
1a, probability density of the incidence *q*_0_. 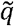 is the upper limit of the *Z*% confidence interval of *q*_0_. This upper limit is termed the maximum plausible incidence. 1b, development of the incidence through time. The incidence growth exponentially. We are interested in estimating the incidence *q*_0_ when our most recent sample takes place. A period *⊿*_0_ earlier a sample was taken ad the incidence at that time was *q*_0_*Z*_1_, a period *⊿*_1_ before that a sample was taken and the incidence was *q*_0_*Z*_2_, etc.

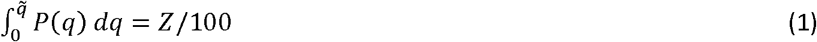

Several methods have been published about repeated sampling of populations to estimate incidence [9, 10, 11]. In these papers the disease incidence is assumed to be constant. In reality with invading pathogens the pathogen population, and equivalently the population of infected hosts will often grow exponentially during the early period of invasion. Following the ideas developed by JAJ Metz [12] several authors have studied the cases of disease freedom and first detection with exponential growth of the number of infected hosts [8, 13, 14, 15]. These authors studied the scenario in which an expert does multiple surveillance rounds, in which they assess a number of plants for the presence/absence of disease, and with a fixed time interval between surveillance rounds. From the data gathered, the maximum plausible incidence, 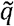, (as defined above) is calculated.

The scenario we study in this paper is that the expert verifies reports from the volunteer and we compare that with the scenario where experts sample for themselves without prior scouting by volunteers. We assume the expert can assess the infection status with certainty for example because they can bring samples into the lab and perform any diagnostics needed. We will derive expressions for the maximum plausible incidence, 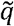. We will compare this maximum plausible incidence when the expert verifies volunteer reports, 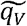, with the scenario where the expert goes into the field and chooses their own hosts to assess for disease, 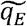. By this comparison, we will be able to quantify the value of volunteer reporting for the early detection of an invader. It is apparent ahead of the analysis that, broadly, the value will depend on the accuracy of the volunteer identification, the extra sampling effort made available by involving volunteers, the cost of expert verification, and the cost of responding to detections. Our purpose is to derive general results which explain how these various quantities combine to determine the value of voluntary surveillance.

In the material and methods section, we describe the generic cases for sampling to establish disease freedom and first detection. These lead to the use of numerical procedures to calculate the maximum plausible incidence. These methods are possible to use by specialists but are of little value to practitioners. Moreover using numerical solutions only it is not easy to get general insight into the effect of parameters on the maximum plausible incidence. We therefore derive a series of approximations that are explicit and can be used by practitioners. The approximations also give insight into the value of volunteer surveillance for early detection of invaders. We will assess the accuracy of the approximations by comparing the maximum plausible incidence calculated from the full model and the approximations.

## MATERIAL AND METHODS

We will use 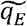 and 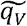 to denote the upper limit of the confidence interval of *q*_0_ for sampling by experts only and for verifying reports of volunteer surveyors, respectively. We use 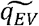 for surveys including both experts sampling on their own and validation of reports of volunteers. In the sections where approximations are compared with exact solutions we will use 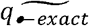 and 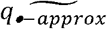, where • can be *E* or *V*, to denote the exact and the approximated upper limit, respectively.

### Preamble

#### 1. The probability for a volunteer surveyor to report a positive host

Disease incidence is denoted by *q*. The probability that a volunteer surveyor observes an infected host to be uninfected, the false negative rate, is *θ*_*fn*_. The probability that the volunteer surveyor observes an uninfected host to be infected, the false positive rate, is *θ*_*fp*_. We denote the uninfected as 0 and the infected as 1. Table 1 summarises the probabilities. The probability for the volunteer surveyor to observe an infected, 1, host is

**Table 1:** Table of the disease status and the observation of infected, 1, and uninfected, 0, hosts. The incidence of disease in the host population is q. θ_fp_ is the false positive rate of the observations. θ_fn_ is the false negative rate of the observations.

|  |  | Disease status |  |
| --- | --- | --- | --- |
| | | 1-$q$ | $q$ |
|  |  | 0 | 1 |
| observation | 0 | $(1-q)(1-\theta_{fp})$ | $q\theta_{fn}$ |
| | 1 | $(1-q)\theta_{fp}$ | $q(1-\theta_{fn})$ |

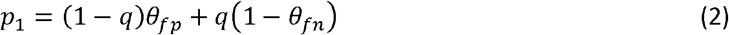

The probability to observe an uninfected, 0, host is

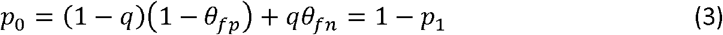

After some rearrangement we see from (2) and (3) that the probability, *k*_1_, to be a false positive given that the volunteer surveyor reports a positive is

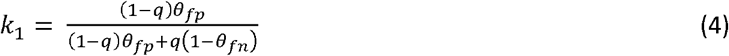

#### 2. Multiple monitoring rounds

We assume that the epidemic is growing exponentially in time with rate *r*. This assumption is reasonable because we are only interested in small values of *q*. The incidence increases as 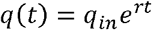, where *q*_in_ is the initial incidence. We want to estimate the incidence at the most recent monitoring round, *q*_0_. At each previous monitoring round the incidence was smaller, see figure 1b. We will number the monitoring rounds starting with 0 for the most recent monitoring round. The time interval between two monitoring rounds is Δ. From the exponential growth we find that *Z*_*i*_ = (*e*^*r*Δ^)^−*i*^ := *λ*^−*i*^, and thus *q*_*i*_ = *λ*^−1^*q*_0_. λ can be interpreted as the multiplication factor of the incidence in a Δ time step, figure 1b.

### Disease freedom sampling

#### 1. Regulatory survey only

In a monitoring programme of K rounds (where the most recent round is round 0 and the first round is round K) the expert samples *N*_*K*_, *N*_*K-1*_, …., *N*_2_, *N*_1_, *N*_0_ hosts. The expert concludes that none of these hosts are positives. We denote the number of true positives in monitoring round *i* buy *Y*_Ni_. When the incidence is *q*_i_ the probability not to find any infected hosts in a sample of size *N*_i_ is (1-*q*_i_)^*N*i^. Therefore, the probability not to find any infected hosts in all *K* monitoring rounds is given by:

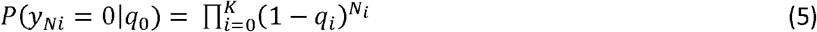

We will use Bayes’ equation to calculate *P*(*q*_0_|*y*_Ni_=0),

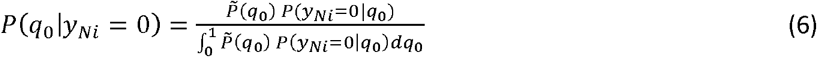

We assume that there is no pre-existing knowledge of the incidence and thus the prior, 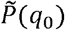, is taken as a uniform density between 0 and 1. This results in

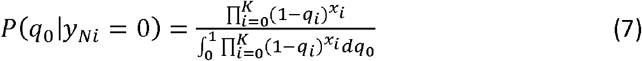

Using equation (1) and (7), and noting that *q*_i_=*λ*^-*i*^*q*_0_, we can now numerically calculate the upper limit of the *Z*% confidence limit of *q*_0_, 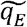. This *q*_0_ is informally called, as discussed above, the maximum plausible incidence.

#### 2. Volunteer surveillance only

The volunteer surveyor reports *x*_*K*_, *x*_*K-1*_, …., *x*_2_, *x*_1_, *x*_0_ hosts assessed as being infected. We denote the number of true positives in monitoring round *i* by *y*_*xi*_. In the absence of disease all hosts reported by the volunteer surveyor are verified by the expert and found not infected, 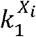. The probability not to find any infected hosts in all *K* monitoring rounds is thus given by:

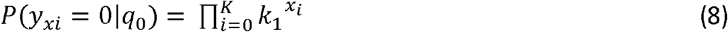

Using Bayes’ equation to calculate *P*(*q*_0_|*y*_*i*_=0) as above we find

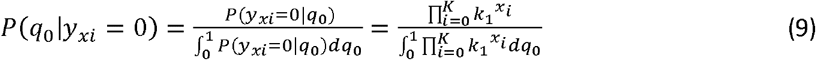

Using equation (1) and (9), and noting that *q*_i_ =*λ*^*-i*^*q*_0_, we can numerically calculate the upper limit of the *Z*% confidence limit of *q*_0_, 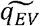.

#### 3. Combined volunteer surveillance and regulatory survey

The volunteer surveyor reports *x*_*K*_, *x*_*K-1*_, …., *x*_2_, *x*_1_, *x*_0_ hosts as infected and all of these are verified by the expert. On top of this the expert samples *N*_*K*_, *N*_*K-1*_, …., *N*_2_, *N*_1_, *N*_0_ hosts themself. In this case

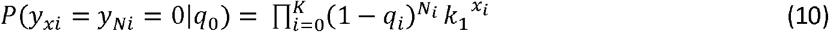

and using Bayes’ equation to calculate *P*(*q*_0_|*y*_i_=0) as above we find

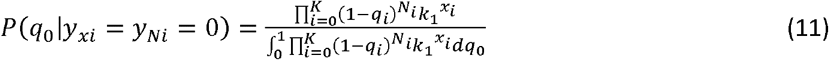

From which we can numerically calculate the upper limit of the *Z*% confidence limit of *q*_0_,, 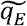.

### First detection

#### 1. Regulatory survey only

Following Parnell et al (2012) the expert samples *N*_*K*_, *N*_*K-1*_, …., *N*_2_, *N*_1_, *N*_0_ hosts. In the survey rounds *K* to 1 none of the sampled hosts is infected, (1-*q*_i_)^*N*i^, *i*∈[*K*,1]. Only in the last round, round *i*=0, one or more sampled hosts turn out to be infected, (1-(1-*q*_0_)^*N*0^). We have:

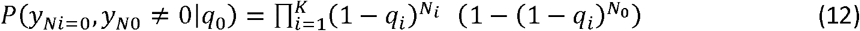

As in the disease freedom case we calculate *P*(*q*_0_ | *y*_*Ni*=0_,*y*_*N*0_ ≠ 0) using Bayes’ equation with a uniform prior and find.

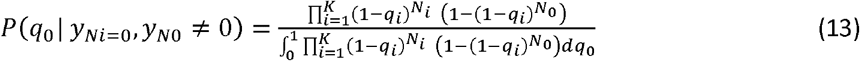

Using equation (1) and (13) we can numerically calculate the upper limit of the *Z*% confidence limit of *q*_0_, 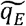.

#### 2. Volunteer surveillance only

The volunteer surveyor again reports *x*_*K*_, *x*_*K-1*_, …., *x*_2_, *x*_1_, *x*_0_ cases. All reported cases in surveillance round *K* to 1 turn out to be not infected after the expert verifies the finds, 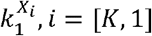. In the surveillance round 0 one or more reported cases are confirmed to be infected after expert verification, 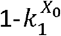. We then get

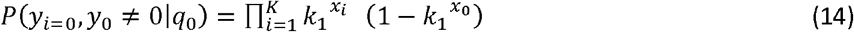

As in the disease freedom case we calculate *P(q*_0_ | *y*_*i*=0_, *y*_0_ ≠ 0) using Bayes’ equation with a uniform prior.

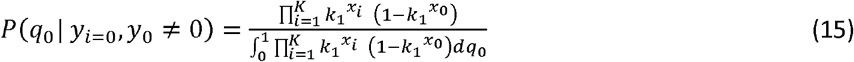

From which we can numerically calculate the upper limit of the *Z*% confidence limit of *q*_0_ 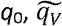.

#### 3. Combined volunteer surveillance and regulatory survey

The volunteer surveyor reports *x*_*K*_, *x*_*K-1*_, …., *x*_2_, *x*_1_, *x*_0_ hosts as infected and all of these are verified by the expert. On top of this the expert samples *N*_*K*_, *N*_*K-1*_, …., *N*_2_, *N*_1_, *N*_0_ hosts him/her self. In survey rounds K to 1 all hosts turn out to be uninfected. In the most recent round, round 0, one or more hosts are found to be infected. We then have:

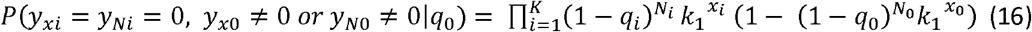

Using Bayes’ equation to calculate *P*(*q*_0_|*y*_*xi*_ = *y*_*Ni*_ = 0, *y*_*x*0_ ≠ 0 *or y*_*N*0_ ≠ 0) as above we find

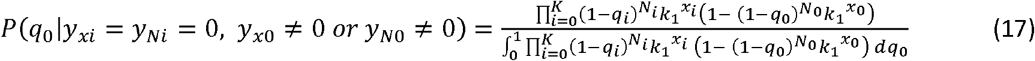

Using equation (1) and (17) we can numerically calculate the upper limit of the Z% confidence limit of *q*_0_,, 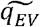.

### Approximations

Equations (7), (9), (11), (13), (15) and (17) can be approximated to give simple expressions for the *Z*% upper limit of the confidence interval for *q*_0_, the maximum plausible incidence. First, we write

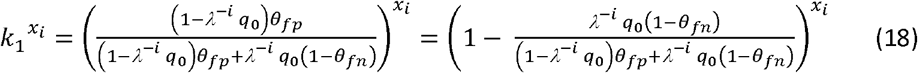

Since we are only interested in small values of *q*_0_ we can write

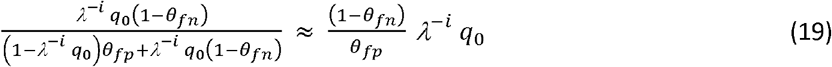

and finally

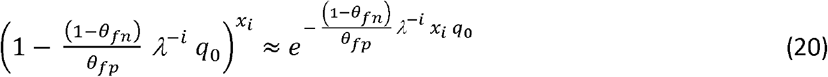

Moreover since we are only interested in small values of *q*_0_ we use 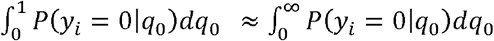 and 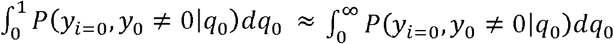.

For the <u>Disease freedom</u> situation, equations (7), (9) and (11), we find that the probability distribution of the incidence, P(*q*_0_), is of exponential form

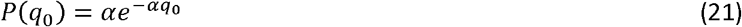

And the upper limit of the *Z*% confidence interval for *q*_0_, is

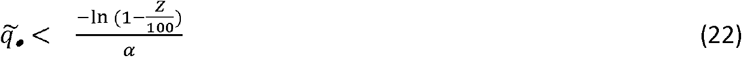

Where 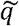 can be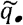 or 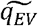 depending on the case under consideration, and α is as defined in Table 2. For the <u>First Detection</u> cases, equations (13), (15) and (17), we find that the probability distribution is a hypo-exponential density.

**Table 2:** Approximations to the probability densities of the incidence of disease in the most recent survey round, *P*(*q*_0_). Densities are given for the case of disease freedom, where all survey rounds return no positive finds, and for first detection where in the most recent survey round one of more positives is found. For the disease freedom cases the right-hand column gives the upper limit of the *Z*% confidence interval for the incidence. For the case of first detection this upper limit is approximated using the z-score.

| DISEASE FREEDOM | Probability density | Max. incidence |
| --- | --- | --- |
| Expert only | $P(q_0) = \left( \sum_{i=0}^K \lambda^{-i} N_i \right) e^{-\left( \sum_{i=0}^K \lambda^{-i} N_i \right) q_0}$ | $\widetilde{q}_E = \frac{-\ln(1-\frac{Z}{100})}{\sum_{i=0}^K \lambda^{-i} N_i}$ |
| Volunteer surveillance only | $P(q_0) = \left( \frac{(1-\theta_{fn})}{\theta_{fp}} \sum_{i=0}^K \lambda^{-i} x_i \right) e^{-\left( \frac{(1-\theta_{fn})}{\theta_{fp}} \sum_{i=0}^K \lambda^{-i} x_i \right) q_0}$ | $\widetilde{q}_V = \frac{\theta_{fp}}{1-\theta_{fn}} \frac{-\ln(1-\frac{Z}{100})}{\sum_{i=0}^K \lambda^{-i} x_i}$ |
| Combined expert sampling and volunteer surveillance | $P(q_0) = C e^{-C q_0}$<br>$C = \sum_{i=0}^K \lambda^{-i} N_i + \frac{1-\theta_{fn}}{\theta_{fp}} \sum_{i=0}^K \lambda^{-i} x_i$ | $\widetilde{q}_{EV} = \frac{-\ln(1-\frac{Z}{100})}{C}$ |

| FIRST DETECTION | Probability density | Max. incidence |
| --- | --- | --- |
| Expert only | $P(q_0) = \frac{1}{\frac{1}{A} - \frac{1}{B}} (e^{-A q_0} - e^{-B q_0})$<br>$A = \sum_{i=1}^K \lambda^{-i} N_i$ and $B = \sum_{i=0}^K \lambda^{-i} N_i$ | $\widetilde{q}_E = \frac{1}{A} + \frac{1}{B} + z \sqrt{\frac{1}{A^2} + \frac{1}{B^2}}$<br>$z$ is the z-score for the standard normal distribution.<br>For the 95% tail $z=1.96$, for the 99% tail $z=2.576$ |
| Volunteer surveillance only | $P(q_0) = \frac{1}{\frac{1}{A} - \frac{1}{B}} (e^{-A q_0} - e^{-B q_0})$<br>$A = \frac{1-\theta_{fn}}{\theta_{fp}} \sum_{i=1}^K \lambda^{-i} x_i$ and $B = \frac{1-\theta_{fn}}{\theta_{fp}} \sum_{i=0}^K \lambda^{-i} x_i$ | $\widetilde{q}_V = \frac{1}{A} + \frac{1}{B} + z \sqrt{\frac{1}{A^2} + \frac{1}{B^2}}$<br>$z$ is the z-score for the standard normal distribution.<br>For the 95% tail $z=1.96$, for the 99% tail $z=2.576$ |
| Combined expert sampling and volunteer surveillance | $P(q_0) = \frac{1}{\frac{1}{A} - \frac{1}{B}} (e^{-A q_0} - e^{-B q_0})$<br>$A = \sum_{i=1}^K \lambda^{-i} N_i + \frac{1-\theta_{fn}}{\theta_{fp}} \sum_{i=1}^K \lambda^{-i} x_i$ and<br>$B = \sum_{i=0}^K \lambda^{-i} N_i + \frac{1-\theta_{fn}}{\theta_{fp}} \sum_{i=0}^K \lambda^{-i} x_i$ | $\widetilde{q}_{EV} = \frac{1}{A} + \frac{1}{B} + z \sqrt{\frac{1}{A^2} + \frac{1}{B^2}}$<br>$z$ is the z-score for the standard normal distribution.<br>For the 95% tail $z=1.96$, for the 99% tail $z=2.576$ |

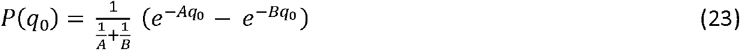

Using equation (1) to calculate the upper limit 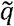 does not give an explicit expression of 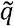 in the model parameters and to obtain an approximation, we appeal to the law of large numbers and the z-score of the standard normal distribution to arrive at an approximation for 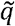. The mean and variance of the hypo-exponential distribution are 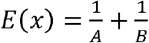 and 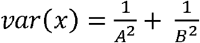, respectively. Now assume that for a large number of samples, the hypo-exponential density can be approximated by a normal density. Then the z-score is

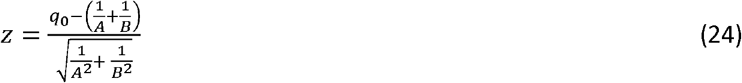

Which for the 95% tail z=1.96, for the 99% tail z=2.576. Solving for 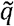 we find

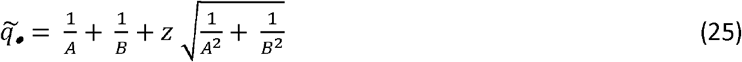

Where 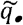. can be 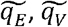, or 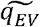 depending on the case under consideration, and A and B are defined in Table 2.

### The accuracy of the approximations

We will calculate for a range of epidemic and surveillance parameters the upper limit of the confidence interval for *q*_0_, for the distributions (7), (9), (11), (13), (15) and (17), 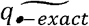, and for their approximating distributions, 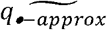, given in Table 2. The relative difference between the two tells us about the accuracy of the approximation. For this analysis we need realistic values of the epidemic growth rate of tree diseases. Table 3 summarises the growth rate of 6 tree diseases, some from natural systems and some from production orchard systems. The graphs to assess the accuracy of the approximations will be made for a pathogen with a large epidemic growth rate, citrus canker, and for one with a small epidemic growth rate, ash dieback.

**Table 3:** Epidemic growth rate of 6 tree diseases of natural forests and agricultural orchards.

| Disease | organism | Mean epidemic growth rate | references |
| --- | --- | --- | --- |
| Ash dieback | *Hymenoscyphus fraxineus* | 0.0024 | Alonso Chavez et al. 2016 |
| Sudden oak death | *Phytophthora ramorum* | 0.0033 | Alonso Chavez et al. 2016 |
| Citrus canker | *Xanthomonas citri* | 0.0184 | Alonso Chavez et al. 2016 |
| Huanglongbing | *Candidatus Liberibacter spp.* | 0.0072 | Alonso Chavez et al. 2016 |
| olive quick decline syndrome | *Xylella fastidiosa* | 0.0122 | Mastin et al in press |
| Pine pitch canker | *Fusarium circinatum* | 0.0019 | Wikler et al 2003; Reynolds et al 2019 |

### Time budgets of the expert and volunteer surveillance

A key constraint in monitoring programmes is money and/or time. It will take experts less time to sample a host themselves than to verify a report from a volunteer surveyor, because of time requirements to transfer the information from the volunteer surveyor to the expert and for the expert to verify that the validation survey is located correctly. So assume the expert has in total *T* time units to do the work. To sample one host themself an expert takes *T*_*N*_ time units, to verify a reported tree it costs *T*_*X*_ time units. Then

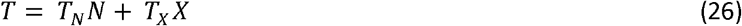

Which is the same as

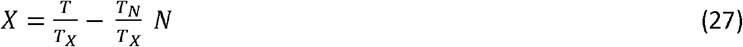

Now consider the probability that the incidence of the disease is smaller than a value *q*\*, which is given by

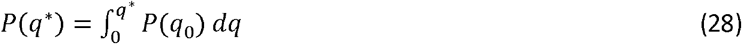

Obviously with the monitoring programme one wants to maximise the probability *P*(*q*\*) that the incidence is below *q*\*, for all values of q*. Equation (28), which is a function of *N* and *X*, allows us to plot contour lines of equal value of *P*(*q*\*) in the *N-X* plane (see figure 5). Superimposing the budget constrained (27) on that plot it is possible to identify the conditions under which it is time effective to verify reports of volunteer surveyors.

## RESULTS

### Approximations

Table 2 summarises the approximations to the upper limit of the *Z*% confidence interval of q_0_, which we termed the maximum plausible incidence. Given that the false-positive and false-negative rates of the volunteer surveyor are known these equations enable us to calculate the maximum plausible incidence both in the case of disease freedom and in the case of first detection. With this information we can address the question of the value for experts to verify reports of volunteer surveyors instead of sampling themselves. If in both cases the expert samples/verifies *N* trees, so *N=x* we see from table 2 that in both the situation where the disease is absent and for the first detection case

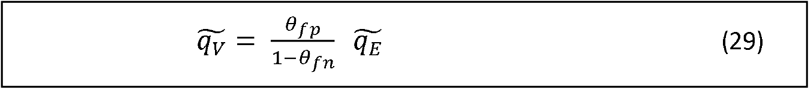

The maximum plausible incidence is a factor 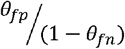 smaller/larger when the experts verify reports of volunteer surveyors than when the experts sample on their own. Figure 2a shows lines of equal value of this factor as a function of the false-positive and the false-negative rate. We note that θ_fp_ is also known as the false positive proportion, FPP, and, 1-θ_fn_ is also known as the true positive proportion, TPP. The FPP/TPP ratio measures the value of volunteer surveillance.

**Figure 2:**
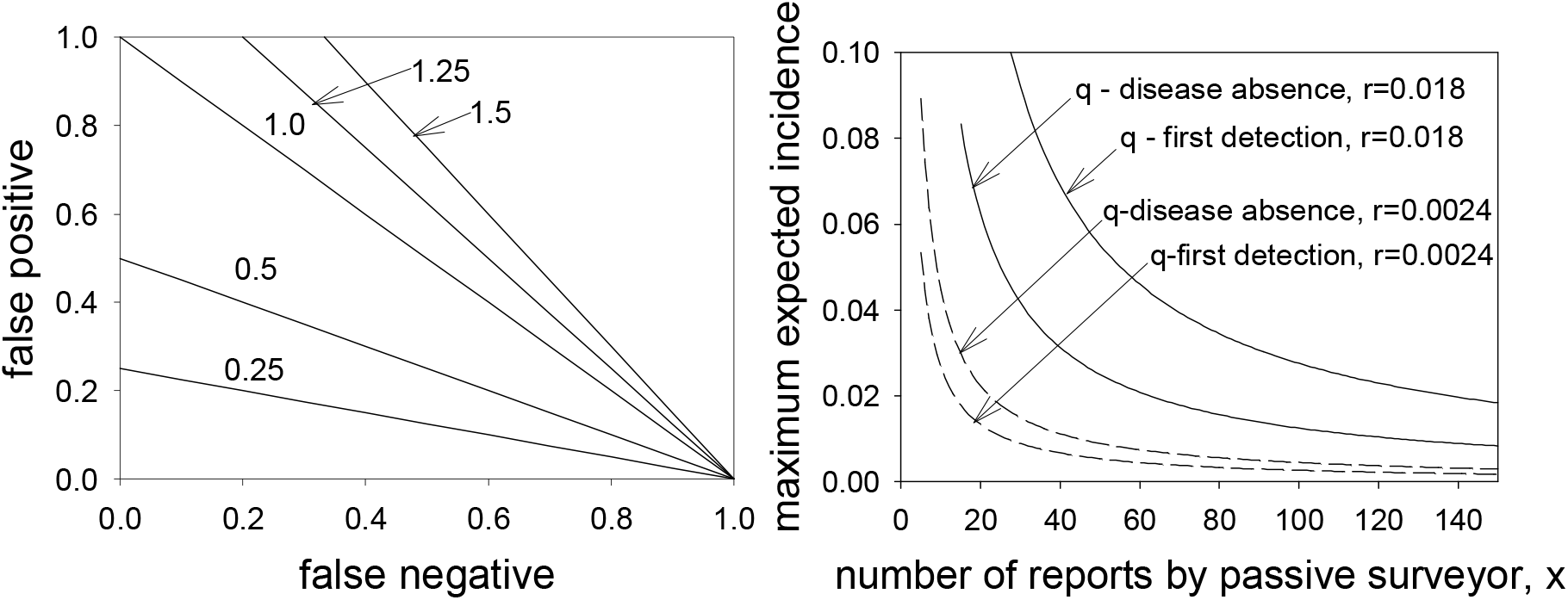
Left-hand panel: Contour lines of 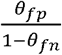 for values ofθ_fn_ and θ_fp_. Right-hand panel: The maximum plausible incidence (i.e. the right hand boundary of the 95% confidence interval for *q*_0_) as function of the number of cases reported by the volunteer surveyor. Drawn lines are for a pathogen with an epidemic growth rate of 0.018 (comparable to Citrus Canker), the dashed line for a pathogen with growth rate 0.0024 (comparable to Ash Dieback). The maximum plausible incidence is shown both for (i) the case where during all monitoring rounds the expert, verifying reports of volunteer surveyors, does not find any host to be infected (disease freedom), and (ii) when in the last monitoring round the expert, verifying reports of the volunteer surveyor, does find an infected host.

### The accuracy of the approximations

The accuracy of the approximation of 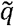 was quantified by

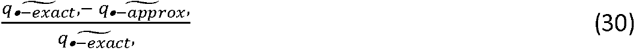

Where 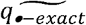 is the upper limit of the *Z*% confidence interval for *q*_0_ in the full model and 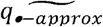 is the upper limit calculated for the approximation. The accuracy of the approximations 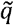 for the experts sampling on their own has been studied [13, 15]. We study the accuracy of the scenario where experts verify the reports of volunteer surveyors only. Figure 3 shows the results of the analysis. Clearly, both the approximation for the disease freedom case and for the first detection case are more accurate for smaller epidemic growth rates. The approximations are however surprisingly accurate. Even for survey intervals of 3 months, for samples larger than 30, the difference between the approximation and the full model is less than 5%. For samples larger than around 15 the difference is less than 10%.

**Figure 3:**
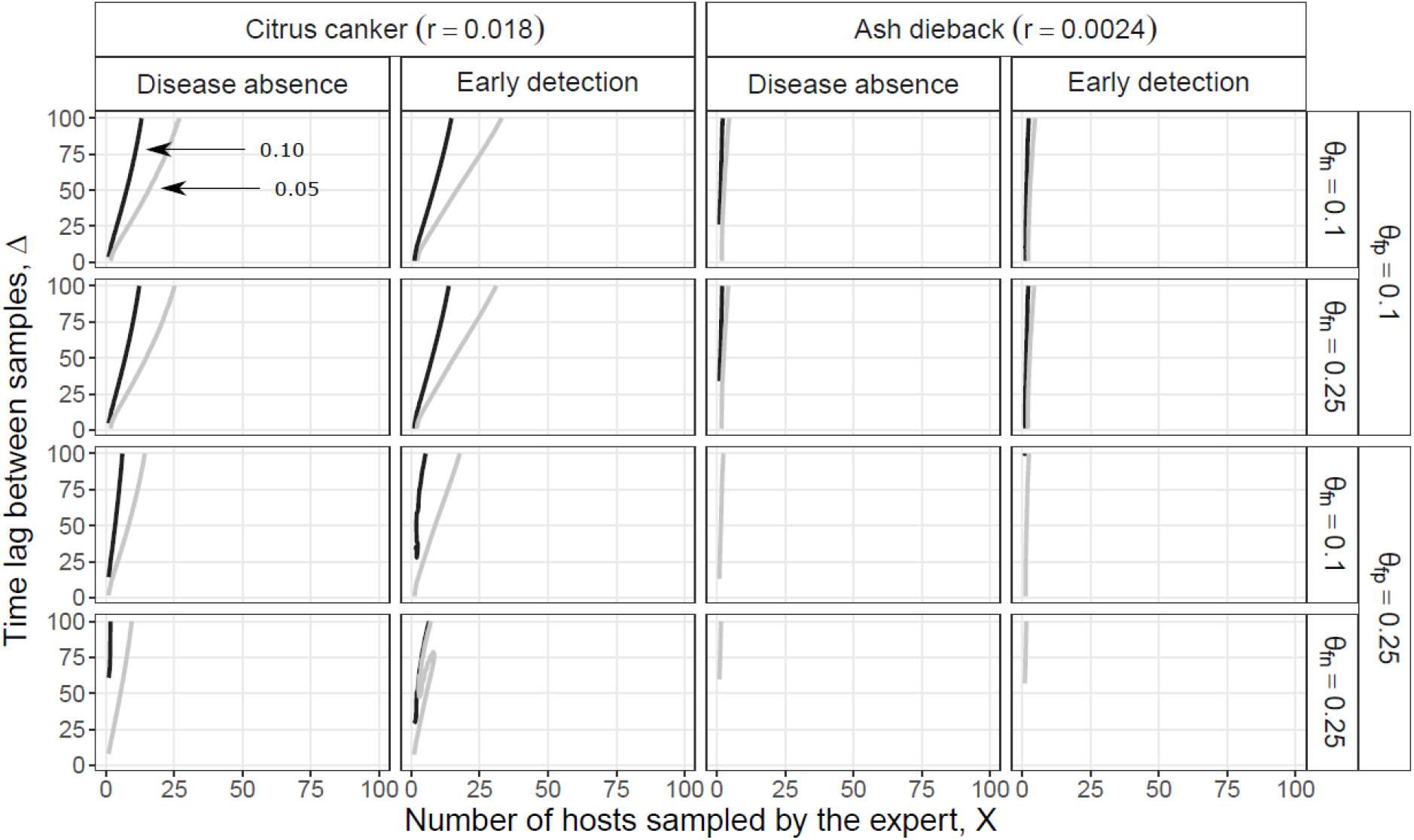
A comparison between the maximum plausible incidence, *q*_0_, as calculated from the approximation and as calculated from the full model. Both disease freedom and first detection is considered for a range of false-positive and false-negative rates for two tree diseases, Citrus Canker and Ash Dieback. On the left-hand side of the black line the value of *q*_0_ calculated from the approximation is less than 10% different from that of the full model. On the left-hand side of the grey line the value calculated from the approximation is less than 5% different from the full model.

### Accuracy of the 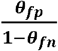 ratio

The 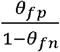 ratio quantifying the gain of involving volunteer surveillance into a programme to detect exotic invaders is derived from the approximations. To see whether this ratio is also a good description of the gain of involving volunteer surveyors when the full models are used, we calculated the upper limit of the confidence intervals of *q*_0_, for the full model of the expert sampling on their own, 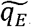 and the full model for the expert verifying reports of the volunteer surveyor,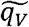 The ratio of these two, 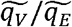, was compared with the 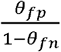 ratio. Figure 4 shows the results of this analysis. As with the accuracy of the approximation of the upper limit of the confidence interval for *q*_0_, the 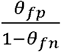 ratio is less than 5% different from 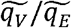 when more than 35 samples are taken. The ratio is less than 10% different from 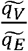 when more than 20 samples are taken.

**Figure 4:**
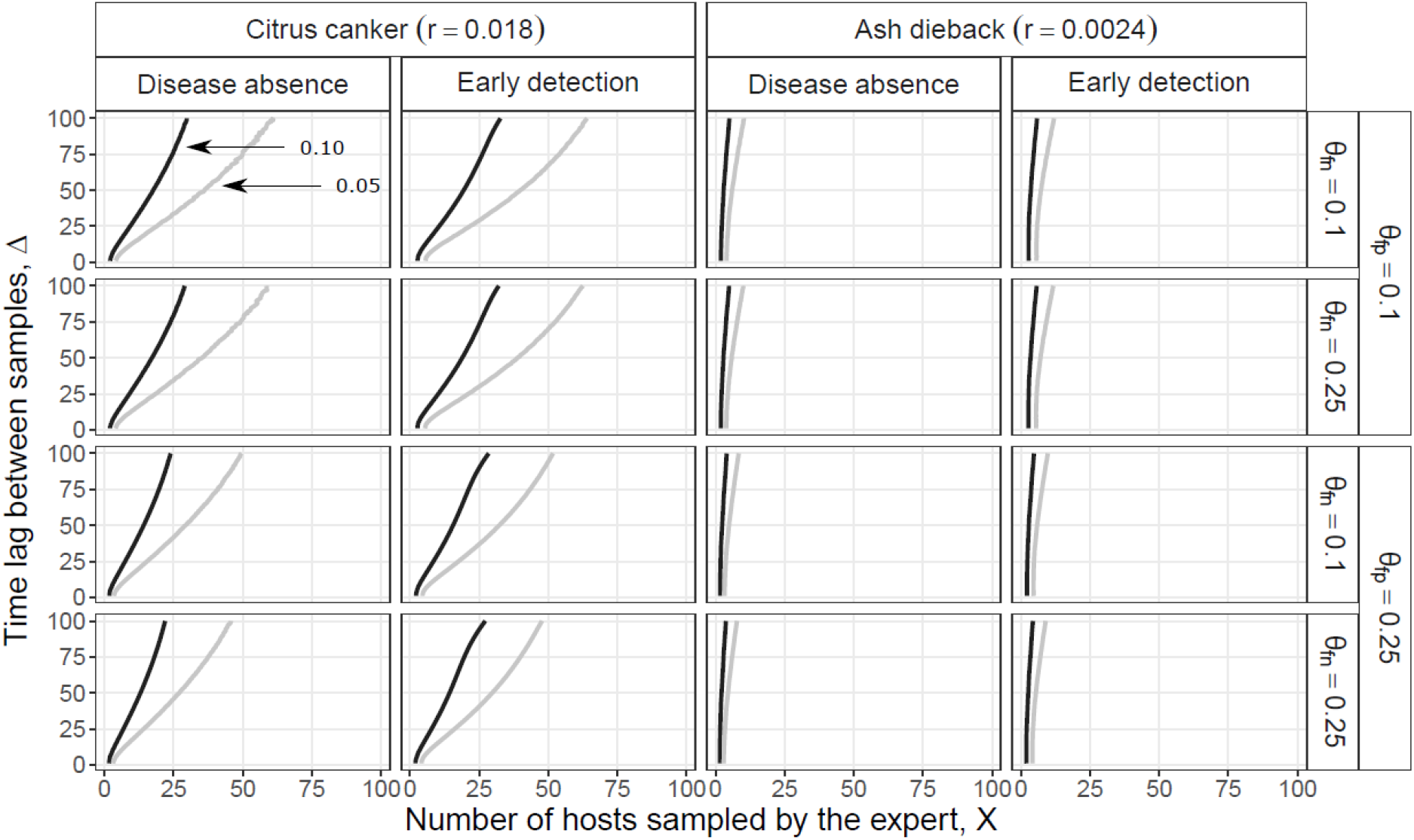
The accuracy of the factor 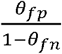. The approximations show that 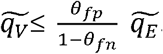. 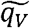 is the maximum plausible incidence when the expert only validates cases reported by the volunteer surveyor to be positive. 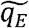 is the to be expected incidence when the expert goes into the field themselves and sample. From the full model we calculate 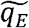 and 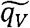. The ratio of these is compared to the factor. 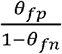 Both disease freedom and first detection is considered for a range of false-positive and false-negative rates for two tree diseases, Citrus Canker and Ash Dieback. On the left-hand side of the black line the value of the factor is less than 10% different from that calculated from the full model. On the left-hand side of the grey line the value of the factor is less than 5% different from that calculated from the full model.

### Difference between disease freedom and first detection

Figure 2b shows the maximum plausible incidence in the case of disease freedom sampling and in the case of first detection for the disease with a small epidemic growth rate and for the disease with a large epidemic growth rate. The figure shows that the estimated incidence in the case of first detection is between 1.5 (for low epidemic growth rate) and 2.5 (for high epidemic growth rate) times the incidence in the case of disease freedom.

### Time budgets of the expert and volunteer surveillance

#### Disease freedom

From (28) we find

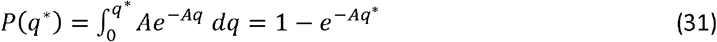

And solving for *X* we get

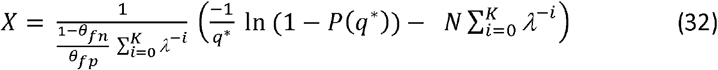

So these are lines with intercept 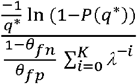 and slope 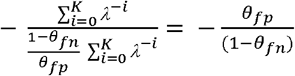

Figure 5 shows both (27) in orange and (32) in blue for different values of *P*. The aim is obviously to maximise *P* for all possible values of q*. From the graphs we conclude that the optimal surveillance programme is to

**Figure 5:**
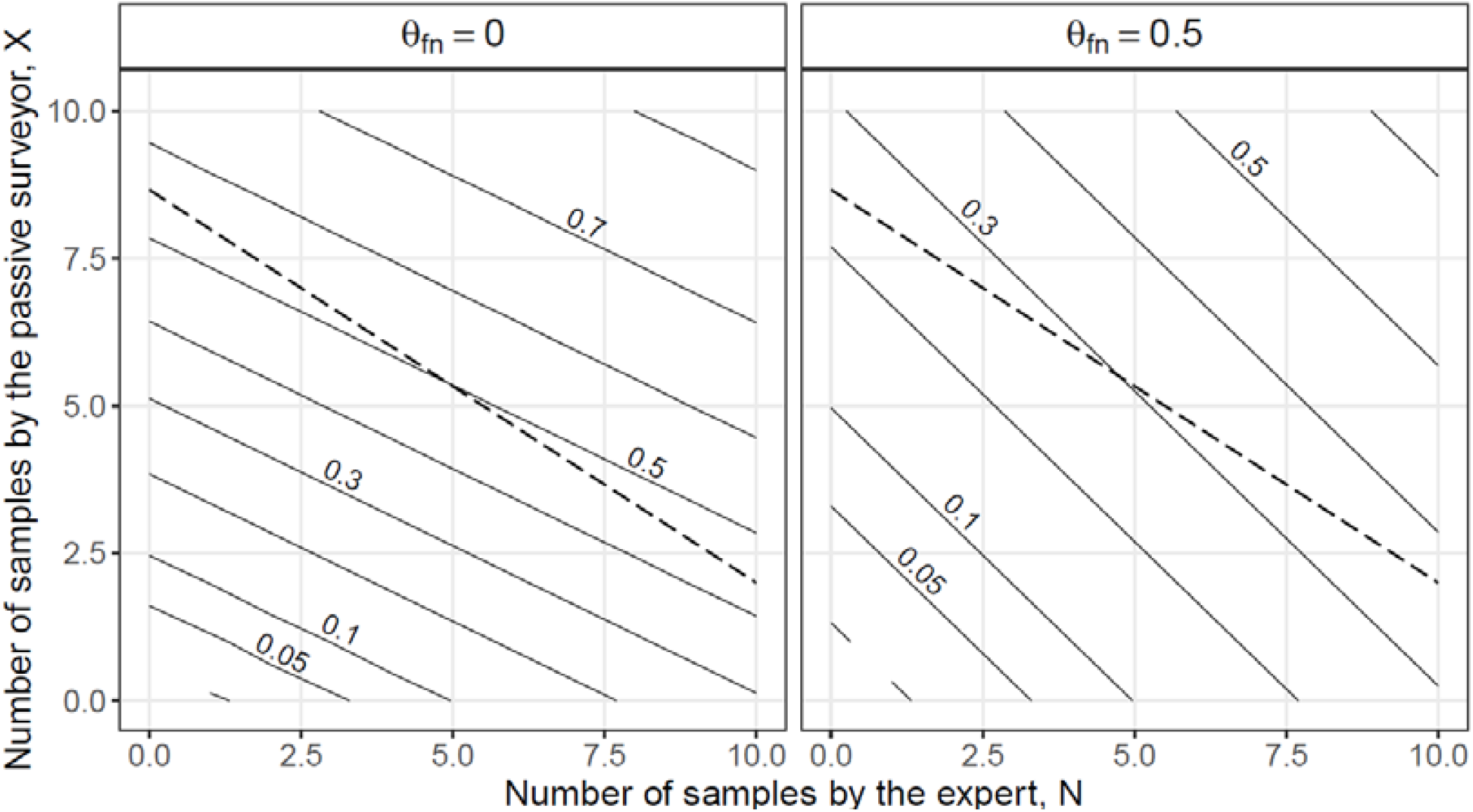
Lines of equal probability that the disease is found before incidence *q*. each drawn line is the contour line for a value of *q*_0_. The hashed line is the contour line for equal total cost of the monitoring programme. The left hand graph shows a case where the optimal monitoring programme consist of experts only verifying the reported cases of the volunteer surveyor. The right-hand panel shows a case where the optimal surveillance programme consist of experts going into the field themselves to sample.

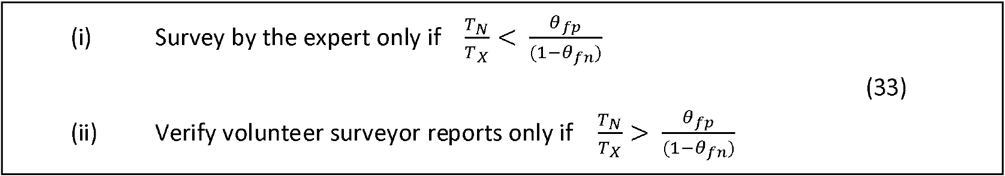

#### First detection

From (28) we get

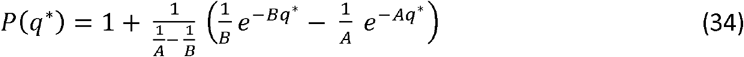

Where

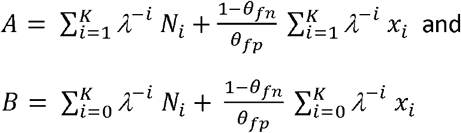

In this case it is not possible to express *X* as function of *N* and model parameters. Contour lines were drawn numerically. An example is given in figure 5. The contour lines for equal P from (34) are virtually indistinguishable from straight lines (supplementary materials *I* gives a large set of graphs showing the generality of this statement). Moreover, the slope of the lines is virtually indistinguishable from 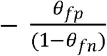. This implies that in practice the same conclusion is reached for the case of first detection as that derived for the disease freedom case.

## DISCUSSION

Volunteer surveillance accumulate potentially valuable datasets for research [5]. False-positive and false-negative observations are, however, a concern about the usefulness of the data, and as discussed in the introduction these rates can be large in some cases and quite small in others. Using statistical techniques, it is possible to estimate false-positive and false-negative rates and correct for them. For example, [16] used citizen science data to create an accurate early warning system for the occurrence of the Asian tiger mosquito in Spain. Brown et al [17] combined observations from volunteer surveyors (forest owners, forest managers) and a regulatory survey to produce a map of Acute Oak Decline in the UK. Cruickshank et al [7] reconstructed occupancy trends of amphibians based on citizen scientists reporting.

These examples have one thing in common, they use the volunteer surveyor data, calculate a measure of the false-positive and false negative rate and in using the data include the measured error rates in the calculation of density and trend. The present case of early detection of invading exotic species is different in that a reported observation of an exotic invader cannot lead automatically to the assumption of the establishment of the invader. The reporting will always need to be verified by an expert. The question then is what the value is of volunteer surveillance reports. Should they be used as a preselection of sites/trees to be visited or is that not an effective use of the expert’s time?

We have shown that the maximum plausible incidence of the disease when volunteer surveillance reports are verified is a factor 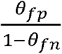 smaller (or larger) than the maximum plausibled incidence when the experts sample on their own. Given that both the false-positive and the false negative rates are small including volunteer surveyor into surveillance programmes can potentially be of great benefit. There is however a possibility that including volunteer surveyors has a negative effect. When the false-negative rate is larger, the factor can be larger than 1 (Figure 2a). It is not entirely clear whether that will happen in practice. If for example the false positive rate is 0.2, as in the amphibian example [7], the false negative rate needs to be close to 0.8 before the 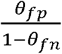 ratio becomes larger than 1, which seems prima facie unlikely. It is much more to be expected that false positive and false negative rates are smaller than 0.5, the equivalent of flipping a coin. In that case the gain from including volunteer surveyors into surveillance programmes for the early detection of exotic invaders will always be positive. This is a useful result since doing better than coin flipping in assigning infected/infested status is the mildest minimum capability criterion one could imagine for this type of activity and performance far in excess of this is likely to be a requirement in any practical situation.

We have developed a range of approximations on the basis of which the maximum plausible incidence can be calculated when the false-positive and false negative rates are known. These equations can be used to determine the maximum plausible incidence from data on the number of hosts verified and the outcome of that verification. The equations can also be used in the development stage where a surveillance programme is set up and one needs to know how many hosts should be assessed to be sure that the maximum plausible incidence is below a pre-set threshold.

For both types of calculations we need an estimate of the epidemic growth rate, *r*, and of the false-negative and false-positive rates for the volunteer surveyor. For invading pathogens, the epidemic growth rate is not normally known. In such cases information on past invasions and/or invasions at other places can be used together with mechanistic insight into the effect of the difference in the environments is likely to have on differences in epidemic growth rate [13,18].

Estimating false-positive and false-negative rates has been done in some cases recently [6, 7, 17]. Deriving such estimates is time consuming and there are therefore not many existing examples. The simplest approach has volunteer surveyors assess hosts that have also been assessed by experts. The expert assessment then can be used as the gold standard and the false-positive and false negative rates of volunteer surveyors estimated. The need for expert assessment is often the most costly part of the exercise. It would be worthwhile to investigate whether a technique to estimate false positive and false negative rates for diagnostic tests, the latent class analysis [19], can be used in this case as well. For that analysis no gold standard is needed. A range of diagnostics is used to assess the disease status of a group of hosts, the technique then both separates the hosts into an uninfected and an infected group as well as estimating the false positive and false negative rates of each of the diagnostics. Having a group of volunteer surveyors assess the disease status of a number of hosts is an essentially comparable process and it is to be expected that latent class analysis could be used to estimate false-positive and false-negative rates for non-expert surveyors.

Several authors [13, 15] have assessed the accuracy of the approximations for the maximum plausible incidence in the cases where the experts sample on their own. Here we have quantified the accuracy for the cases where experts verify reports of volunteer surveyors (figure 3). In both cases it was shown that for the range of epidemic growth rates observed in reality, (i.e. values of *r* between 0.002 and 0.02 per day) the approximations deviated less than 5% form the full model when the number of samples assessed was larger than 50. The approximations deviated from the full model by less than 10% when the number of samples exceeded 25. We conclude that the approximations are accurate enough to be useful in a practical situation where other stochastic factors are likely to add uncertainty to the detection process. Parnell and Bourhis arrived at very similar conclusions for the approximations to methods where the experts sample on their own.

Finally, we investigated whether verifying volunteer surveyor reports is time effective or whether the expert going into the field on his/her own to sample hosts is the more time effective method. We have shown a very simple rule for when reports of volunteer surveyors should be verified. This rules say that if the ratio of the time an expert needs to sample a host him/herself and the time needed to verify a report of a volunteer surveyor and is larger than the factor 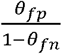, the most time effective method is to dedicate experts’ time only to verification of the work of volunteer surveyors. This gives a clear criterion for when verifying reports by volunteer surveyors should be included in the development of regulatory surveillance programmes. In case volunteer surveillance and expert sampling is combined our results also show how to estimate the maximum plausible incidence.

## REFERENCES

1. Ferguson N, Donnelly C, Anderson R. 2001 Transmission intensity and impact of control policies on the foot and mouth epidemic in Great Britain. Nature 413, 542–548. (http://dx.doi.org/10.1038/35097116).

2. Morath S, Fielding N, Tilbury C, Jones B. 2015 Oriental Chestnut Gall Wasp News of a recent unwelcome discovery and how ‘citizen science’ can play an important role in surveying and identification. Quarterly Journal of Forestry 109, 253–258. (http://dx.doi.org/10.5197/j.2044-0588.2020.041.034).

3. Perez-Sierra A, Gorton C, Kalantarzadeh M, Sancisi-Frey S, Brown A, Hendy S. 2015 First report of shot blight caused by Siroccus tsugae on atlantic cedar (Cedrus atlantica) in Britain. Disease Reports Published Online:17 Sep 2015 (https://doi.org/10.1094/PDIS-04-15-0378-PDN).

4. Dickinson JL, Zuckerberg B, Bonter DN. 2010 Citizen Science as an Ecological Research Tool: Challenges and Benefits. Annual Review of Ecology, Evolution and Systematics 41, 149–172. (http://doi.org/10.1146/annurev-ecolsys-102209-144636).

5. Encarnacao J, Teodosio MA, Morais P. 2021 Citizen science and biological invasions: a review. Frontiers in Environmental Science 8:602980. (doi: 10.3389/fenvs.2020.602980).

6. Falk S, Foster G, Comont R, Conroy J, Bostock H, Salisbury A, Kilbey D, Bennett J, Smith B. 2019 Evaluating the ability of citizen scientists to identify bumblebee (Bombus) species. PLoS ONE 14(6), e0218614. (https://doi.org/10.1371/journal.pone.0218614).

7. Cruickshank SS, Bühler C, Schmidt BR. 2019 Quantifying data quality in a citizen science monitoring program: False negatives, false positives and occupancy trends. Conservation Science and Practice. 2019;e54. (https://doi.org/10.1111/csp2.54).

8. Bourhis Y, Gottwald TR, van den Bosch F. 2018 Translating surveillance data into incidence estimates. Philosophical Transactions of the Royal Society B Biological Sciences 374(1776):20180262. (doi: 10.1098/rstb.2018.0262).

9. Cameron AR, Baldock FC. 1998 Two-stage sampling in surveys to substantiate freedom from disease. Prev. Vet. Med. 34 (1), 19–30. (doi: 10.1016/S0167-5877(97)00073-1)..

10. Cannon RM. 2002 Demonstrating disease freedom-combining confidence levels. Prev. Vet. Med. 52 (3), 227–249. (doi: 10.1016/S0167-5877(01)00262-8).

11. Coulston JW, Koch FH, Smith WD, Sapio FJ. 2008 Invasive forest pest surveillance: survey development and reliability. Can. J. For. Res. 38 (9), 2422–2433. (doi: 10.1139/X08-076).

12. Metz JAJ, Wedel W,Angulo AF. 1983 Discovering an Epidemic before it has Reached a Certain Level of Prevalence. Biometrics 39, 765–770.

13. Parnell S, Gottwald TR, Cunniffe NJ, Alonso Chavez V, van den Bosch F. 2015 Early detection surveillance for an emerging plant pathogen: a rule of thumb to predict prevalence at first discovery. Proc. R. Soc. B 282: 20151478. http://dx.doi.org/10.1098/rspb.2015.1478

14. Mastin A, van den Bosch F, Gottwald TR, Alonso Chavez V, Parnell SR. 2017 A method of determining where to target surveillance efforts in heterogeneous epidemiological systems. PLoS Comput Biol. 28, 13(8):e1005712. (https://doi.org/10.1371/journal.pcbi.1005712)

15. Bourhis Y, Gottwald TR, Lopez-Ruiz FJ, Patarapuwadol S and van den Bosch F. 2019 Sampling for disease absence-deriving informed monitoring from epidemic traits. Journal of Theoretical Biology 461, 8–16. (doi: 10.1016/j.jtbi.2018.10.038).

16. Palmer JRB, Oltra A, Collantes F, Delgado JA, Lucientes J, Delacour S, Bengoa M, Eritja R, Bartumeus F. 2017 Citizen science provides a reliable and scalable tool to track disease-carrying mosquitoes. Nature Communications 8, 916 (doi:10.1038/s41467-017-00914-9).

17. Brown N, van den Bosch F, Parnell S, Denman S. 2017 Integrating regulatory surveys and citizen science to map outbreaks of forest diseases: acute oak decline in England and Wales. Proc. R. Soc. B 284: 20170547. (http://dx.doi.org/10.1098/rspb.2017.0547).

18. Gottwald TR. 2010 Current epidemiological understanding of citrus huanglongbing. Annual Review of Phytopathology 48, 19–139. (http://doi.org/10.1146/annurev-phyto-073009-114418).

19. Turechek WW, Webster CG, Duan J, Roberts PD, Kousik CS, Adkins S. 2013 The use of latent class analysis to estimate the sensitivities and specificities of diagnostic tests for Squash vein yellowing virus in cucurbit species when there is no gold standard. Phytopathology 103, 1243–1251. (doi: 10.1094/PHYTO-03-13-0071-R).

20. Alonso Chavez V, Parnell S, Van Den Bosch F. 2016 Monitoring invasive pathogens in plant nurseries for early-detection and to minimise the probability of escape. Journal of Theoretical Biology 407, 290–302. (doi:10.1016/j.jtbi.2016.07.041).

21. Mastin A, van den Bosch F, Bourhis Y, Parnell S. 2022 Out of sight: Surveillance strategies for emerging vectored plant pathogens. Scientific Reports, in press. BioRxiv repository (doi: https://doi.org/10.1101/2022.01.21.477248).

22. Wikler K, Storer AJ, Newman W, Gordon TR, Wood DL. 2003 The dynamics of an introduced pathogen in a native Monterey pine (Pinus radiate) forest. Forest Ecology and Management 179, 209-221. (doiI:10.1016/S0378-1127(02)00524-8).

23. Reynolds GJ, Gordon TR, McRoberts N. 2019 Projecting monterery pine (Pinus radiate) populations over time in the presence of various representations of pitch canker disease, caused by Fusarium circinatum. Forest 10, 437. (doi:10.3390/f10050437).

