## Supplementary materials 1 for "The value of volunteer surveillance for the early detection of biological invaders"

Contour plots of equal probability P(q*), as defined in equation (28) as function of the number of hosts sampled by the expert, N, and the number of reports of the volunteer surveyor, X. Plots are presented for two epidemic growth rates, r=0.018 and r=0.0024, and for several values of the false-positive and false-negative rates. The results show that the contour lines are virtually indistinguishable from straight lines and that the slope is virtually indistinguishable from θ_fp_/(1-θ_fn_).

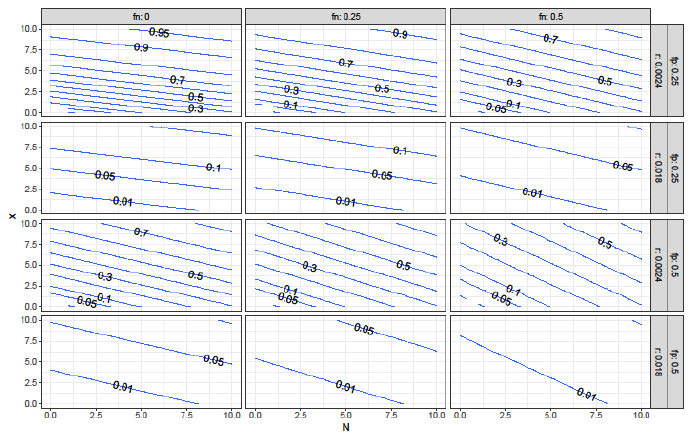
